# TPR1, a novel rifampin derivative demonstrates efficacy alone and in combination with doxycycline against the NIAID Category A Priority Pathogen *F. tularensis*

**DOI:** 10.1101/2021.02.12.431005

**Authors:** Jason E. Cummings, Keaton W. Slayden, Richard A. Slayden

**Author notes:** Author to whom correspondence should be addressed. **RAS:** Department of Microbiology, Immunology and Pathology, Colorado State University, Fort Collins, CO 80523-2025.

## Abstract

**Objectives:** *Francisella tularensis* is a highly virulent and contagious gram-negative intracellular bacterium that causes the disease tularemia in mammals and is classified as a Category A priority pathogen. The high infectivity, difficulty of obtaining a durable cure of disseminated disease and the emergence of drug resistance warrants investigation of new drugs that can be used alone and in combination with existing standard of care clinical drugs.

**Methods:** We utilized a systematic analysis of antibacterial potency, extent of dissemination by analysis of bacterial burden in a secondary vital organ, and survival rates to assess the efficacy of a novel rifampicin derivative, TPR1. The efficacy of TPR1 was evaluated alone and in combination with the standard of care drug, doxycycline, against Type A *F. tularensis* Schu S4 using a lethal pulmonary model of infection in mice.

**Results:** TPR1 has an MIC value range of 0.125-4 mg/L against reference laboratory strain SchuS4 and a panel of clinical strains. TPR1 alone reduced the bacterial burden in the lungs and spleen at 40 mg/kg and 80 mg/kg, and no antagonism was observed when co-administered with doxycycline. Dosing at 40 mg/kg doxycycline reduced the bacterial burden by 1 Log_10_ CFU in the lungs and 4 Log_10_ CFU spleen in comparison to untreated controls. Co-administration of TPR1 and doxycycline demonstrated efficacy upon treatment withdrawal after 4 days of treatment, and 100% survival.

**Conclusions:** Significantly, TPR1 demonstrated efficacy when delivered alone and in combination with doxycycline, which provides compelling evidence of a superior treatment strategy that would normally rely on a single chemotherapeutic for efficacy. In addition, this work substantiates the use of rifampicin derivatives as a platform for the development of novel treatments to other bacterial agents in addition to tularemia.

## Introduction

*Francisella tularensis* is a highly virulent and contagious Gram-negative intracellular bacterium that causes the disease tularemia in mammals and is often fatal when acquired by inhalation.^1, 2^ In addition to being an emerging difficult to treat pathogen, NIAID has classified *F. tularensis* as a Category A priority pathogen. Tularemia is distributed primarily in the northern hemisphere including North America, Europe, Japan, China, the Middle East and Russia.^3^ Tularemia outbreaks are commonly reported in these endemic regions and typically associated with contaminated water or soil, wild and domestic animals, and common arthropod vectors^3^. Thus, there is an important and continuing need for new treatments for tularemia.

Streptomycin and gentamicin are considered the first-line clinical drugs, and other treatment regimens for *F. tularensis* infections utilize rifampicin, doxycycline and ciprofloxacin.^4^ The use traditional chemotherapeutics including streptomycin and gentamicin are often limited because of toxic side effects, and neither can be orally administered. Despite the availability of drugs such as the aminoglycosides, macrolides, chloramphenicol and fluoroquinolones, and tetracyclines, relapse of disease is found in 10% of the cases and can result in a mortality as high as 40%. ^4, 5^ Fluoroquinolones have shown promise in a limited number of cases; however, clinical application is limited and their role in treating severe disease is unknown. In animal studies, gatifloxacin, moxifloxacin, and ciprofloxacin prevented disease during the treatment period, but significant failure rates occurred after the withdrawal of therapy. ^3^ Since existing drugs have limited potency and efficacy, and it is unknown whether drug-resistant organisms might be used in an intentional release, it is prudent to develop novel chemotherapeutics with proven modes of action and reduced side effects that can be co-administered with standard of care (SoC) drugs to treat acute infections and prevent relapse of disease.

Doxycycline and rifampicin have been shown to have activity against *F. tularensis*. ^6^ However, the activity of these drugs against *F. tularensis* has been reported to not achieve a durable cure, and the emergence of drug resistance has been observed. ^5, 7^ *Ex vivo* studies have demonstrated that rifampicin penetrates well within eukaryotic cells and demonstrates activity against intracellular *F. tularensis*, but development of resistance and frequent disease relapse is observed upon withdrawal of drug treatment. ^5, 6^ Similarly, doxycycline has been shown to be efficacious against *F. tularensis* but has been associated with frequent clinical disease relapses. ^1^ There are numerous drug treatment guidelines for *F. tularensis*, with the recommendation to administer streptomycin and gentamicin for at least 10-14 days as well as tetracyclines and chloramphenicol for at least 14-21 days. ^3, 6, 8^ In severe cases, two drug combinations are recommended to prevent relapse and drug resistance.

An historically successful approach in drug discovery is repurposing or to utilize a known pharmacophore as the foundation for derivatization to create a new drug. ^9–11^ The identification and use of a derivative of a widely used broad-spectrum promises to improve management of emerging pathogens, such as *F. tularensis*. The bacterial DNA-dependent RNA polymerase is a widely exploited drug target in bacteria making rifampicin an ideal pharmacophore as a foundation for new derivatives. In this study, we assessed the use of a novel rifampin-derivative, TPR1, alone and in combination with doxycycline to reduce bacterial burden, prevent bacterial dissemination and relapse, and increase survival. *In vitro* results show TPR1 being equally active against a diverse panel of *F. tularensis* strains that included clinical isolates and reference laboratory strains. The minimum inhibitory concentration (MIC) value of TPR1 was 0.125-4 mg/L, which is comparable to current SoC drugs used to treat infections caused by *F. tularensis*. ^6^ TPR1 also demonstrated efficacy in a mouse model of *F. tularensis* infection when administered alone at 80 mg/kg and in combination with doxycycline at sub-efficacious doses of 40 mg/kg as determined by a reduction in bacterial burden in the lungs and spleen with no observable disease relapse. This substantiates the use of the rifampicin derivative, TPR1, alone or in combination with doxycycline for the treatment of infections caused by *F. tularensis* that will provide a durable cure without relapse or the emergence of drug resistance associated with single drug treatments.

## Results and discussion

### TPR1 is a potent inhibitor of F. tularensis with low cytotoxicity

TPR1 was screened against a diverse panel of *F. tularensis* strains consisting of laboratory and clinical strains that are considered representative of the drug susceptibility spectrum associated with clinical infections. TPR1 has a minimal inhibitory concentration range of 0.125-4 mg/L against the laboratory reference strain and a panel of clinical strains with various susceptibilities to SoC drugs (Table 1). This MIC value is characteristic of clinical drugs with bactericidal activity against *F. tularensis*, which is within the range for a drug to have efficacy. Most significantly, TPR1 demonstrated potency comparable to current clinically used drugs to treat *F. tularensis* infections. For example, the MICs determined for various clinical strains of *F. tularensis* with streptomycin and gentamicin fall into a range of 2-4 mg/L and 1-2 mg/L, respectively. To assess the potential safety window of TPR1, cytotoxicity was evaluated in HepG2 cells. The LC_50_ for TPR1 was determined to be >128 mg/L, which is comparable to other clinical drugs and represents a significant difference in the selective index (SI) between tissue toxicity and bactericidal dose (Table 1).

**Table 1.** Minimal inhibitory concentration (MIC) and HepG2 cell line cytotoxicity (LD_50_)

| MIC mg/L | TPR1 | Rifampicin | Doxycycline | Gentamicin |
| --- | --- | --- | --- | --- |
| *F. tularensis* Schu S4 Submaster Cell Bank<br>(BEI NR-10492) | 4 | 0.5 | 1 | 1 |
| *F. tularensis* Schu S4 FSC237 (BEI NR-643) | 2 | 0.25 | 2 | 1 |
| *F. tularensis* WY96-3418 (BEI NR-644) | 0.125 | 0.0625 | 0.125 | 0.25 |
| *F. tularensis* MA00-2987 (BEI NR-645) | 4 | 0.5 | 2 | 1 |
| *F. tularensis* KY99-3387 (BEI NR-647) | 4 | 0.5 | 2 | 1 |
| *F. tularensis* OR96-0246 (BEI NR-648) | 2 | 0.25 | 0.5 | 0.5 |
| HepG2 Cytotoxicity | >128 | N/A | N/A | N/A |

### TPR1 demonstrates efficacy and controls dissemination

The efficacy of TPR1 was determined in the standard *F. tularensis* murine model of infection. For these studies, mice were infected with *F. tularensis* Schu4 and administered TPR1 once a day for 5 days beginning day one, and monitored for a total of 28 days for morbidity, mortality, and disease relapse. Mice from each treatment group, untreated and control group were selected to be euthanized, and the bacterial loads in lung and spleen was determined by plating and enumeration of CFU. Untreated mice generally displayed 6 log_10_ CFU in the lung at day 4 (Figure 1A) and 7 log_10_ CFU in the spleen (Figure 1B). The bacterial burdens detected in the lungs and spleen were consistent with the observed rapid weight loss and 0% survival rate with a median survival of 4 days (Figure 1CD). The efficacy of TPR1 was assessed at 40 mg/kg and at 80 mg/kg. Both levels of TPR1 dosing demonstrated a reduction in bacterial burden in the lungs and spleen compared to untreated controls. When delivered at 40 mg/kg, TPR1 reduced bacterial growth in the lungs and no bacteria were detected in the spleen (p<0.01) (Figure 1AB). The observed partial efficacy of TPR1 delivered at 40 mg/kg is consistent with the observed delayed weight loss and increased median survival of 8 days (Figure 1CD). Durable cure was achieved when TPR1 was delivered at 80 mg/kg based on no detectable bacteria in the lungs or spleen, reversible weight loss and 100% survival until the study endpoint of 7 x the median survival of untreated controls (Figure 1). This is consistent with the observed differences in efficacy and demonstrates correlations with median survival, bacterial burden in the lungs, control of dissemination, and growth in the spleen during treatment.

**Figure 1.**
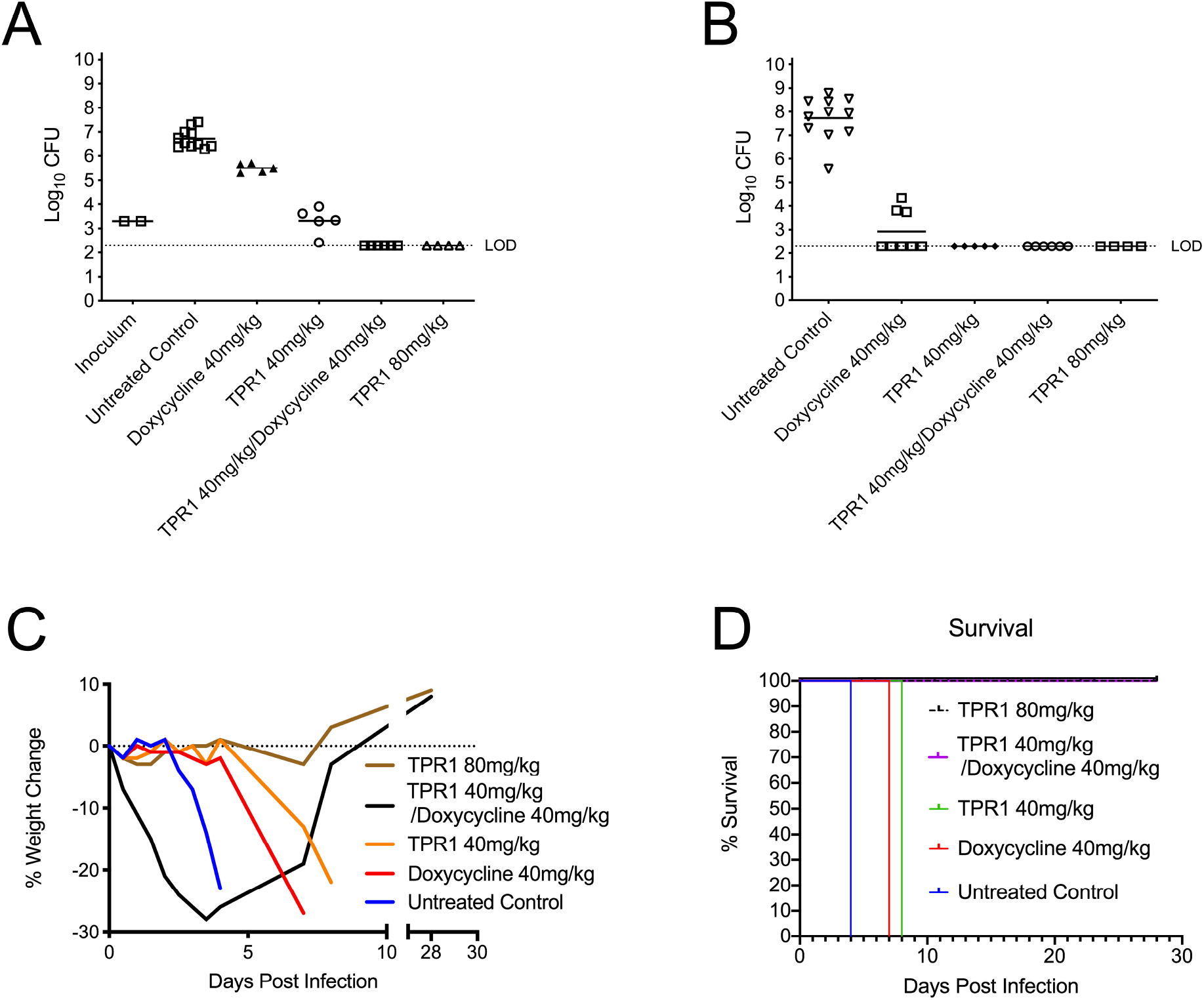
Efficacy in murine model of tularemia. Bacterial burdens in the lung and spleen during treatment with TPR1. *F. tularensis* colony forming units (Log_10_ CFU/mL) recovered from the lungs (A) and spleen (B) at day 4 of treatment. (C) Percent change in animal body weight, and (D) Survival of mice treated with TPR1 or doxycycline, and TPR1 doxycycline co-administration.

### TPR1 and doxycycline co-administration at subtherapeutic doses demonstrate efficacy

TPR1 was assessed as a co-therapeutic with the SoC drug doxycycline to determine its use as a new agent in a combination therapy regimen. Dosing at 40 mg/kg doxycycline reduced the bacterial burden by ∼1 Log_10_ CFU in the lungs and ∼4 Log_10_ CFU spleen in comparison to untreated controls (Figure 1AB), which demonstrates only partial efficacy at this dose. Co-administration of sub-efficacious doses of TPR1 and doxycycline demonstrated efficacy upon treatment withdrawal after 4 days of treatment, and 100% survival until the study endpoint of 28 days, equivalent to ∼7 times the mean time of death of untreated controls (Figure 1CD). Mice that reached study endpoint were shown to have undetectable bacteria in both the lung and spleen. Importantly, this demonstrates that the combination of TPR1 and doxycycline have an additive effect and are capable of obtaining a “durable” cure even when co-administered at sub-efficacious doses.

In this study, we demonstrated that TPR1 possesses antibacterial activity against a panel of *F. tularensis* strains that includes clinical strains with diverse susceptibilities to current SoC drugs. TPR1 was also shown to have efficacy in the murine model of tularemia. The improved efficacy of TPR1 alone and in combination with the SoC drug, doxycycline, in the acute model of tularemia with no observable resistance clearly indicates that this rifampicin derivative has excellent potential as a next-generation chemotherapeutic against tularemia.

## Methods

### Minimal inhibitory concentration of TPR1

To assess activity of TPR1 (supplied by Palisades Therapeutics/Pop Test Oncology LLC) against *F. tularensis*, minimal inhibitory concentration (MIC) assays were performed using the *F. tularensis* laboratory reference strain Schu4 and a panel of clinical strains. MICs were determined *via* CLSI guidelines M07/M45 as previously described. ^12^ *F. tularensis* was grown to mid-log and diluted to provide an inoculum of ∼10^5^ cells/mL when dispensed into a 96-well microtiter drug plate. Compounds in the drug plates were tested at 2-fold serial concentrations from 0.0156-32 μg/ml in triplicate. Optical density readings were obtained after 18-20 hours of incubation at 37°C and MIC defined as the first concentration to exhibit no growth.

### Determination of Cytotoxicity

HepG2 cells were seeded in 96 well plates at 10,000 cells per well in Eagle’s Minimum Essential Medium (ATCC) supplemented with 10% (v/v) fetal bovine serum (cEMEM). Drug plates were prepared with 1:2 fold dilutions starting with 128 mg/L and ending with 0.0625 mg/L in cEMEM. Drug media was added to washed HepG2 cells and incubated at 37°C with 5% CO_2_ for 24hrs. Media was removed and replaced with cEMEM without phenol red. 0.01mL of 12mM methylethiazole tetrazolium (MTT Sigma) was added to the plates and cells incubated for an additional 4 hours at 37°C with 5% CO_2_. 0.1mL detergent solution (0.1g/mL SDS in 0.01M HCl) was added to each well and plates incubated for an additional 4 hours at 37°C with 5% CO2. Absorbance of plates were measured at 570nm to determine percent growth reduction over the concentration series. Values were plotted using Prism software and IC50 values (lethal concentration causing 50% loss of cell viability) determined by linear regression analysis.

### Efficacy and survival studies

Seven to nine week-old female BALB/c mice were purchased from Charles River Laboratories (Jackson Laboratories, Bar Harbor, Maine). All mice were housed in micro-isolator cages in the laboratory animal resources facility or in the Infectious Diseases Research Complex BSL-3 facility at Colorado State University and were provided sterile water and food ad libitum. All research involving animals was conducted in accordance with animal care and use guidelines and animal protocols were approved by the Animal Care and Use Committee at Colorado State University. Mice were anesthetized with a mixture of ketamine/xylezine to achieve 100mg/kg ketamine and 10mg/kg xylezine in a volume of 0.1mL per mouse delivered i.p.. Mice were infected with 100 CFU Schu4 delivered intranasally as previously described. ^2,12, 13^ Mice were monitored for morbidity and mortality twice daily.

TPR1 was dissolved in DMSO to a final concentration of 160mg/mL, resulting in a dosing stock that was further diluted in sterile water dropwise while mixing to a final concentration of 8mg/mL. Treatments were delivered at 100uL per 20g bodyweight P.O. for 40mg/kg or with 200ul per 20g bodyweight P.O. for 80mg/kg. Doxycycline (160mg) was dissolved into 20mL PBS and filter sterilized. Mice were treated with 100uL per 20g bodyweight P.O..

The study design consisted of seven experimental groups: Group 1: untreated control (5% DMSO/Water P.O. B.I.D); Group 2: doxycycline (PBS 40 mg/kg P.O. B.I.D.); Group 3: TPR1 (5% DMSO/Water 40mg/kg P.O. B.I.D.); Group 4: doxycycline::TPR co-treatment (TPR1 5% DMSO/Water 40 mg/kg P.O. B.I.D., doxycycline – PBS 40 mg/kg P.O. B.I.D.); Group 5: TPR1 (5% DMSO/Water 80mg/kg P.O. B.I.D.). Experimental endpoints were 96 hours post-infection, based on the mean time untreated animals become moribund, and 28 days post-infection, representing 7 times the moribund mean time to assess disease relapse. At each 96-hour timepoints, representative animals from each group were sacrificed, and bacterial burden in the lungs and spleen were determined by plating and enumeration of colony forming units (CFUs). The remaining animals in each group were monitored until moribund or the study endpoint if appropriate, and bacterial burden was assessed in vital organs.

## Acknowledgement

We gratefully acknowledge Julie Starkey for assisting with editing and for scientific input, and Dr. Clinton Dawson and Dr. Nurul Islam for scientific input.

## Ethics Statement

Use of vertebrate animals at Colorado State University is conducted under AAALAC approval OLAW number A3572-01 under file with NIH. Animals are housed in ABL-3 facility under supervision by full-time veterinarians per American Veterinary Medical Association guidelines.

## Transparency declarations

All other authors: none to declare.

## Author contributions

The determination of MIC, toxicity and efficacy studies was performed at Colorado State University by J. E. C. with support from K. W. S. Study design and writing was performed by J. E. C. and R. A. S.

